# Tracking changes in birds’ interaction milieu

**DOI:** 10.1101/2025.02.14.638220

**Authors:** Stanislas Rigal, Vincent Devictor, Vasilis Dakos

**Affiliations:** TETIS, Université de Montpellier, AgroParisTech, Cirad, CNRS, INRAE, Montpellier, France; ISEM, Université de Montpellier, CNRS, IRD, EPHE, Montpellier, France

**Keywords:** Biotic homogenisation, bird community, Convergent Cross Mapping, co-occurrence, network, species associations

## Abstract

As biodiversity is declining, the dynamics of species interactions is a growing conservation concern. However, estimating and monitoring explicit species interactions across large spatial and temporal scales remain challenging. An alternative and yet under-explored approach is to track whether and how the interaction milieu, defined as the background of all realised interactions, is changing in space and time. Here, we assess changes in the interaction milieu of common bird communities in France. We estimate associated species pairs using spatial and temporal information for 109 species monitored across 1,969 sites during 17 years. We validate the ecological significance of associated species pairs by testing the relationship between the propensity to be associated and species functional proximity or shared habitat preference. We reconstruct association networks for these intra-guild bird communities and track temporal changes in network layout in terms of size, density of links, modularity and degree distribution. We show that, beyond changes usually documented based on species numbers and abundances, the interaction milieu is also changing non-randomly. Communities become smaller with a similar relative number of associations that becomes unevenly distributed through time. These structural changes vary among bird communities according to their habitat and may impact community functioning and how communities can cope with global change.

## 1. Introduction

Vertebrate communities have been experiencing rapid and profound changes in recent decades, mainly documented through variations in species numbers, abundances, or distributions (Kampichler et al. 2012, Koerner et al. 2017). Changes affecting vertebrate communities have been also analysed through the lens of species ecological traits, e.g. by using community weighted indices that have highlighted a general pattern of homogenisation (Clavel et al. 2011, Davey et al. 2012, Li et al. 2018). However, addressing changes in the structure of the interaction networks underlining these communities have been more challenging (McGill et al. 2006). Any change in sign, strength or number of interactions has an effect on the network structure of a community and can strongly affect community dynamics (Lyons et al. 2016) and functioning (e.g. productivity, efficiency of resource use) (Fontaine and Thébault 2015, Besson et al. 2019). Modifications affecting networks with pair-wise interactions of a given type such as plant-pollinator or predator-prey networks have been related to specific consequences for the conservation of these communities (Tylianakis et al. 2010). However, isolating specific types of interactions remains challenging in many communities, in particular in large-scale studies (Barner et al. 2016). The *interaction milieu* was therefore proposed as a concept to refer to the resultant biotic interaction background to which an individual is exposed when arriving in a community (McGill et al. 2006). This notion thus enables to focus on community changes beyond species loss or turnover when addressing conservation issues in communities for which explicit interactions are out of reach.

Bird communities, in particular, have been extensively studied in particular in terms of changes in species composition at large scale, highlighting a visible and rapid effect of anthropogenic pressures leading to a biotic and functional homogenisation (Kampichler et al. 2012, Lindström et al. 2013, Gaüzère et al. 2015). More locally changes in bird communities have also been assessed through dedicated network analyses based on food webs (Lurgi et al. 2012), plant-frugivore networks (Ramos-Robles et al. 2016), competition for nests (Orchan et al. 2013) and specific types of bird communities such as mixed-species flocks (Goodale et al. 2015, Zou et al. 2018). The observed shifts in species composition indicate that the interaction milieu in common bird communities is also likely to be altered, although the extent and direction of this change may depend on the community’s environment (Rigal et al. 2022). In particular, in the context of the sharp decline in the abundance of birds breeding in open environments (Reif and Hanzelka 2020, Rigal and Knape 2024, Gómez-Catasús et al. 2025), we can expect specific changes in the interaction milieu of these communities (e.g. in farmland, open environment) compared to other environments (e.g. forest, urban environment).

The interaction milieu does not aim at mimicking a definite network of interactions, but rather corresponds to the background of interaction. The distribution of species ecological traits has been initially proposed to assess the interaction milieu in plant communities (McGill et al. 2006) and later extended to microbial and soil communities (Jaillard et al. 2018, Davison et al. 2024). For mobile species such as birds, large-scale monitoring data have been used to infer key elements of the interaction milieu through species association based on community assembly processes: species with similar functional traits or habitats may either tend to repulse (or even exclude) each other because of resource competition, under the limiting similarity hypothesis (Martin and Bonier 2018) or attract each other through social information exchange and facilitation processes (Gil et al. 2017, Seppänen et al. 2007). While the nature of interaction net effect that emerge as the result of several interacting processes between species cannot be defined in communities at large scale (Barner et al. 2016), estimating the interaction milieu becomes more accessible through increasing availability of monitoring data with large spatial and temporal resolutions (Ovaskainen et al. 2016), providing that species’ associations are correctly described.

Species associations have been estimated using spatial co-occurrences of species (HilleRisLambers et al. 2012, Letten et al. 2017), disentangling the effect of the biotic filter (*i.e.* interactions) from other filters (*e.g*. habitat filtering) on co-occurrence patterns (Morueta-Holme et al. 2016, Elo et al. 2021) or in combination with interactions described in the literature (Lurgi et al. 2012). Species associations have also been obtained from species time-series using dynamical system analyses (Ushio et al. 2018, Barraquand et al. 2021). Using such associations as meaningful proxies of ecological interactions has been criticised, due to numerous unchecked non-biotic processes (*e.g.* dispersal capacity, habitat, phylogeography, Moran effet) (Sander et al. 2017, Freilich et al. 2018, Dallas et al. 2019, Blanchet et al. 2020, Cordero and Jackson 2021). However, spatial co-occurrences correspond to an apparent equilibrium resulting from large-scale processes (i.e. biotic and abiotic filters) (Rigal et al. 2022), while fluctuations in species time-series reflect an non-equilibrium inside community composition (Ushio et al. 2018). Estimating the interaction milieu in bird communities can therefore benefit from combining spatial and temporal information on bird species abundance.

Pairwise species associations per se are not the primary focus of a community analysis based on the interaction milieu (McGill et al. 2006) but can be used to quantify, via network metrics, the structure of the interaction milieu (Kaiser-Bunbury and Blüthgen 2015, Lau et al. 2017). A network with species as nodes and association as edges between nodes, can be analysed at network, intermediate (i.e. by sub-groups) or node levels (Lau et al. 2017). For instance, at network level, the structure can be defined by the number of nodes, number of edges and connectance (i.e. density of edges). Network structure can be analysed at intermediate level, e.g. by motifs (i.e. recurrent patterns), average path length (i.e. average number of steps between all possible pairs of nodes) and modularity (i.e. extent of network division into sub-groups). At node or species level, the network structure can be studied using, for instance, the degree (i.e. the number of edges by nodes), the centrality (i.e. how influential is a node) or the partnership diversity (i.e. preference for nodes in a network to attach to other similar nodes) (Delmas et al. 2019). The interpretation of each metric on its own is neither relevant nor generalisable, since a change in the index may be due to changes in the size of the network (Heleno et al. 2012), be correlated with the other indicators (Delmas et al. 2019) and depend on the type of interaction studied (Tylianakis et al. 2010).

In this study, we seek to understand the changes in the interaction milieu that accompany the changes in species composition over the last two decades in France. In addition to the overall changes in the interaction milieu of common bird communities, we expect that there are also variations in the changes of interaction milieu across habitats, as bird communities are not exposed to the same pressures and changes in species composition in the different habitats. To do so, we used the French Breeding Bird Survey from 2001 to 2017 to estimate potential associated species pairs using spatial co-occurrences to account for large-scale community composition filters (Fig. 1a), and we refined the selection of associated species pairs using species time-series (Fig. 1b). We then assessed the relevance of the associated species pairs to infer the biotic background of interaction using species ecological traits (Fig. 1c). Lastly, we tracked the temporal changes in the interaction milieu using network metrics (Fig. 1d) and analyse the consequences, for the conservation of bird communities, of changes in the interaction milieu that accompany changes in species composition.

**Figure 1:**
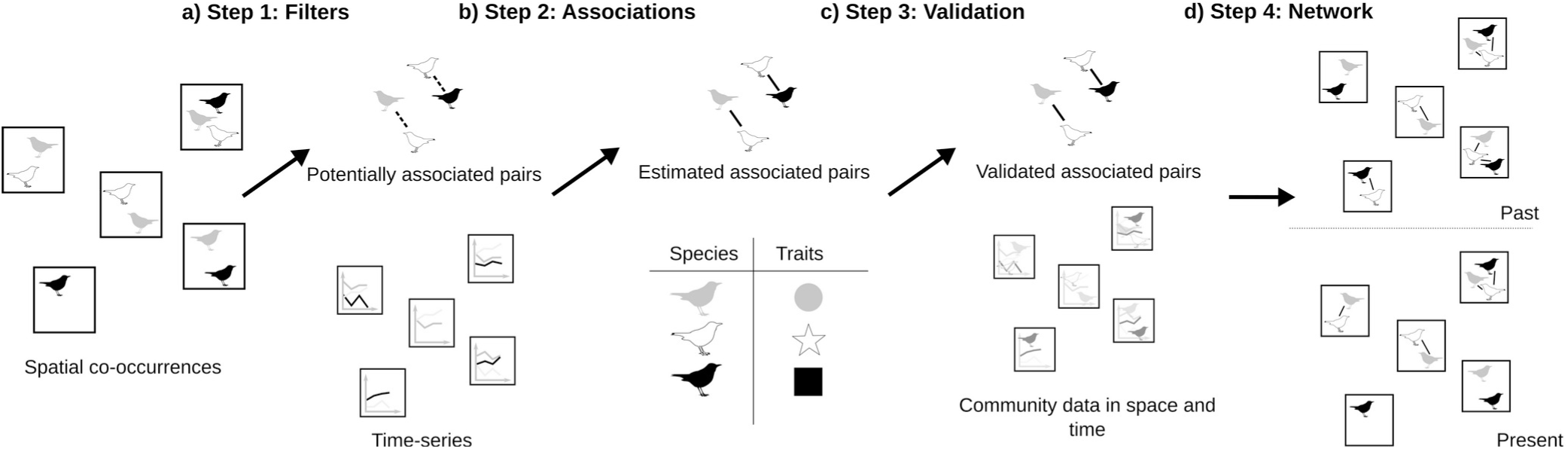
Framework of the estimation of changes in the interaction milieu of bird communities from large-scale spatio-temporal data. a) In a first step, the effects of large-scale processes on species co-occurrences are removed using community apparent equilibrium, to estimate the potentially associated species pairs. b) In a second step, the estimation of associated species pairs is refined by removing the effects of non-biotic local processes on the non-equilibrium of community composition. c) The ecological relevance of the estimated associations is tested using the ecological hypothesis on community assembly and species traits. d) Community composition in space and time are combined with associated species pairs to compute network metrics quantifying the interaction milieu and track its changes.

## 2. Material and methods

### 2.1 Spatial data and time-series of birds

We used the French Breeding Bird Survey (FBBS) based on a standardised protocol to monitor the abundance of common bird species. 2,514 sites of 2×2 km were survey in mainland France between 2001 and 2017 (Jiguet et al. 2012). In each site, the abundances of breeding bird species were counted on 10 sampling points (during 5 min, 1 to 4 h after sunrise) by volunteer experienced ornithologists, twice a year between 1^st^ of April and 15^th^ of June from 2001 to 2017. Each site represented a species assemblage changing in abundance and composition over time. Habitats were described for each sampling point in 19 types (supplementary material 1). The data provided by the French National History Museum (MNHN) were abundances corrected for the difference in species detectability. Among the 242 recorded species, we selected the most abundant species representing 99% of the total abundance in the dataset to avoid over-representation of rare species for which the FBBS is not well designed. After removing rare species and sites only monitored once, our dataset is composed of 1,969 sites and 109 species.

### 2.2 Potentially associated species pairs

We used spatial co-occurrence to estimate the species pairs potentially associated (Fig. 1a) by removing large-scale non-interaction filters (historical dispersal, neutral dispersal and habitat filtering) following Rigal et al. (2022). We measured the partial correlations between species pairs using their co-occurrence profile. We accounted for historical and habitat filtering by estimating potentially associated pairs for each type of habitat and each biogeographic region (see additional details in supplementary material 1). We limited the influence of random processes and uncontrolled habitat filtering using a null model by resampling species in a site. This provided 1000 random co-occurrence datasets obtained by keeping constant the total number of individuals in a given sampling point, and assuming that the probability for a species to occur in a given sampling point was proportional to its frequency in the dataset. We calculated the standardised effect size (SES) of the observed partial correlations compared to the partial correlations from the random datasets. Significant associated pairs were based on p-values (<0.05) obtained from the rank of the observed partial correlations in the distribution of random partial correlations (Morueta-Holme et al. 2016).

### 2.3 Species associations

For each potentially associated species pair in a given habitat and a given biogeographic region, we determined which species pairs were significantly associated (Fig. 1b) based on the relationship between species time-series. We used the multispatial Convergent Cross-Mapping (CCM) method (R package *multispatialCCM*) (Clark et al. 2015), an extension of the CCM method (Sugihara et al. 2012). In CCM, the time-series of two species are linked if we can predict the dynamics of species X using the time-series of species Y, *i.e.* if the attractor of X (defined as the set of states reconstructed from the original and lagged time-series of X) can be estimated from the attractor of Y. If so, species X has a signature into the dynamics of species Y, meaning that species X is affecting species Y (see detailed in supplementary material 2 and Tsonis et al. 2018). CCM resulted in a correlation value *ρ*, depicting the correlation between the two attractors, associated to a p-value for significance.

We focused on sites monitored during at least six consecutive years to maximise the number of sites and with long enough time-series in order to assess significant temporal associations between species. For each species, we reconstructed its time-series at a given site using its recorded abundance, set to zero when the species was not recorded during a monitoring session of the site. Obtaining pseudo-time-series (*i.e.* the times-series reconstructed by placing side by side data from multiple sites) longer than 500 time steps (following the empirical example from Clark et al. 2015) requires to select the 32,491 combinations of species pairs, habitats and biogeographic regions that match this criteria among the 89,270 potentially associated species pairs in given habitat and biogeographic region obtained from spatial co-occurrences. We use 300 replicates for the bootstrap routine due to computational time. The multispatial CCM provides species pairs for which the time-series of one species was related to the time-series of the other. As we aimed at estimating the interaction milieu rather than analysing pairwise associations, we did not account for the direction of the relationship, i.e. if one or both directions was significant, the pair of species was considered to be associated.

### 2.4 Testing ecological relevance of species associations

We investigated whether the associated species pairs were relevant to assess the biotic background of interaction (Fig. 1c) based on trait-based assembly theory and specific expected interactions. According to trait-based assembly theory, species that share similar functional traits, are phylogenetically clustered or with significant habitat overlap, are expected to interact more strongly (Fontaine and Thébault 2015, Mönkkönen et al. 2017). This stronger interaction may be either positive through social information exchange and facilitation processes (Gil et al. 2017), or negative because of resource competition under the limiting similarity hypothesis (Abrams 1983), which should result in a funnel pattern when species interactions are combined with trait-related distances (Mönkkönen et al. 2017). We tested whether the associations obtained from previous steps are in line with these expectations using two approaches. In a first test based on distances between species characteristics, we computed functional, phylogenetic and niche overlap distances (see relationship and complementarity between distances in supplementary material 3) and relate them to the probability of a species pair to be associated. In a second test that is specifically focused on expected interactions, we analysed whether species pairs with a similar diet or nest requirement are more prone to be associated, while controlling for their preferred habitat.

#### 2.4.1 Functional distance

Functional distance was estimated using a life-history trait dataset for the birds of the Palaearctic (Storchová and Hořák 2018). We computed the Generalized Functional Diversity (GFD) following (Mouchet et al. 2008) to generate a pairwise functional distance between all pairs of species (Mouchet et al. 2010). This method calculated all the combinations of the clustering algorithms from a functional distance matrix and provides the best consensus tree, *i.e.* the one for which the dissimilarity with the original distance matrix is the lowest according to the 2-norm goodness-of-fit index (Mérigot et al. 2010). We constructed the initial functional distance matrix using the Gower’s distance as the trait matrix contained both qualitative and quantitative data (Gower 1971). The best consensus dendrogram was built using the Unweighted Pair Group Method using arithmetic Average algorithm (UPGMA) (2-norm=0.03, threshold=3.49). From this dendrogram, we computed the cophenetic functional distance (Sokal and Rohlf 1962) between species, using the 242 initial species to assess the functional distances in the complete functional space of the common bird species.

#### 2.4.2 Phylogenetic distance

Phylogenetic trees of the 242 bird species were obtained from birdtree.org (Jetz et al. 2014). We estimated the consensus tree from 10,000 trees using the *ape* R package (Paradis et al. 2019). We calculated the phylogenetic distance as the cophenetic distance between each pair of species in the consensus tree.

#### 2.4.3 Niche overlap distance

We computed niche overlap for each species pair as the overlap between the frequency distribution of each species within the 19 classes of habitat. To do so, we used the habitat information for each sampling point and we calculated the frequency distribution of each species over the 19 classes of habitats in the whole dataset. We then computed the χ^2^-distance, widely used to compare ecological assemblages (Ben-Shahar and Skinner 1988, Flower et al. 1997), to estimate the distance between each species habitat profile. The χ^2^-distance characterises the proximity in habitat distribution profiles independently of species abundance. We defined the habitat niche overlap distance (*NOD*) using the similarity between habitat profiles as follows (Eq. 1 and see details on NOD in supplementary material 3):

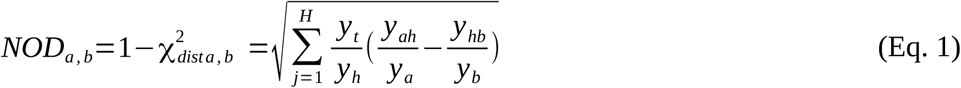

with *NOD_a,b_*the niche overlap distance between species *a* and *b*, *χ*^2^_*dist a, b*_ the χ^2^-distance between habitat profiles of species *a* and *b*, *H* the number of habitats, *y_t_* the total abundance of species *a* and *b* in all the habitats, *y_h_* the abundance of species *a* and *b* in habitat *h*, *y_ah_* the abundance of species *a* in *h*, *y_a_* the total abundance of species *a*, *y_hb_*the abundance of species *b* in *h* and *y_b_* the total abundance of species *b*.

#### 2.4.4 Test with functional, phylogenetic and niche overlap distances

To test the ecological relevance of the associated species pair, we performed a binary logistic regression using Generalised Linear Models (GLM). The response variable (*A*) was the presence or absence of association for each species pair (a species pair is considered to be associated if this has been the case in at least one habitat and one biogeographic region), with a binomial distribution with parameter *p* and a logit link function *g*. The explanatory variables were scaled functional distance (*FD*), phylogenetic distance (*PD*) and niche overlap distance (*NOD*) (FD and PD being slightly correlated but not with NOD, see supplementary material 3) for species pair *i* (Eq. 2):

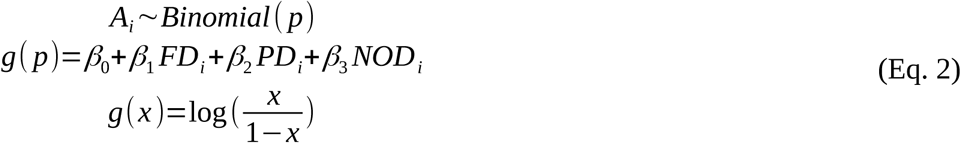

#### 2.4.5 Test with expected interactions for food and nest location

Using the life-history trait dataset for the birds of the Palaearctic (Storchová and Hořák 2018), we recorded for each species pair whether the two species used the same nest type among the five recorded types (*closed arboreal; on ground directly; in tussock very close to ground but not directly on ground; hole in tree, bank, ground, crevice; open arboreal cup in bush, tree, on cliff ledge*) and had the same diet pattern during breeding season based on nine diet descriptors (*at least 10% of diet composed of small plants; fruits; grains and seeds; arthropods; other invertebrates; fish; other vertebrates; carrions; diet composed of similar amount of plants and animals*). To assess the effect of these share traits while controlling for the habitat requirement of the species pair, we recorded whether the two species of the pair prefer the same habitat among the 19 classes, according to their occurrence frequency in each habitat class.

A second GLM was performed in which the binomial response variable (*A*) was the presence or absence of association for each species pair and the explanatory variables were the presence or absence of a share diet (*Diet*), the presence or absence of a shared nest type (*Nest*) and the presence or absence of a similar preferred habitat (*Habitat*) for species pair *i* (Eq. 3).

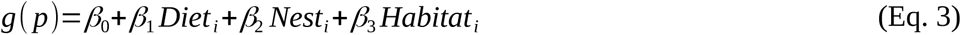

### 2.5 Analysis of the interaction milieu

To quantify the changes in the interaction milieu, we focus on the number of species, the number of associations (Heleno et al. 2012, Kaiser-Bunbury and Blüthgen 2015) and a set of relevant metrics at network, intermediate and node levels. At the network level, we use connectance, a standard metric whose mathematical relationship with other network metrics is well known and which is therefore important for analysing other metrics. The relationship between connectance and perturbations is not consistent, but independently of the type of interactions, connectance increase has been linked to greater species generalism (Tylianakis et al. 2010). At the intermediate level, we use modularity which has been shown as a suitable metric for conservation issues: high modularity restrains perturbations and local extinction within a module (Stouffer and Bascompte 2011). At the node levels, we use evenness of degree distribution that has been related to higher robustness as it decreases the consequences of a local extinction for the remaining species and tend to be lower in the context of degraded habitats (Kaiser-Bunbury and Blüthgen 2015, Fisogni et al. 2021).

#### 2.5.1 Network metrics

We reconstructed a local network (Fig. 1d) at each site and for each year using the species composition (species as nodes) and the associated species pairs (associations as edges). We calculated the following metrics from each network *j*: network size *s* (i.e. number of species), number of associations *a*, connectance *c* (i.e. density of associations in the network), modularity *m* (i.e. intensity of network division into sub-groups) and evenness of degree distribution *e* (i.e. evenness of probability distribution of the number of associations by species). Connectance is the ratio between the observed number of associations *a* and the potential number of associations *a_max_* (Landi et al. 2018) (directly linked to the number of species pairs 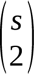, Eq. 4):

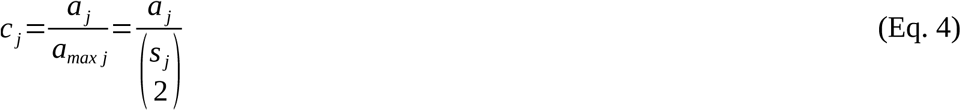

Modularity (Eq. 5) was computed using the R package *rnetcarto* (Doulcier and Stouffer 2015) which enables fast modularity computation (required for the analysis of the amount of networks in this study). It is based on the *rgraph* library and the simulated Annealing method (Guimerà and Nunes Amaral 2005).

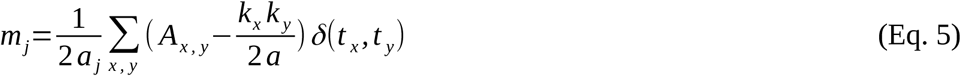

with *a_j_*the number of associations in network *j*, *A_x,y_* the element of the adjacency matrix, *k* the degree and *t* the type to which species *x* or *y* belongs and the Kroenecker delta function defined as equal to 1 when *t_x_ = t_y_* and 0 when *t_x_ ≠ t_y_*.

The evenness of the degree distribution was calculated using the Shannon entropy of the degree distribution (Blüthgen et al. 2008) (Eq. 6):

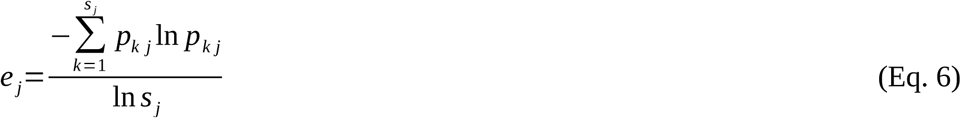

with *p_k_*the frequency of degree *k*.

To explore potential differences in changes of the interaction milieu across habitats, we also conducted the computation of network metrics for the four main habitat classes: farmland, forest, urban and natural open land. We grouped the 19 habitats provided at each site and each year by the volunteers conducting the monitoring into the four habitat classes (see supplementary material 1) and we attributed to each local community its main habitat class to separate between farmland, forest, urban and natural open land communities.

#### 2.5.2 Changes in the interaction milieu through time

We calculated the yearly change in the interaction milieu by estimating the effect of years *Y* (set as an explanatory variable) on each network metric, while explicitly modelling spatial auto-correlation between sites using a Generalised Additive Mixed Model (GAMM) (Eq. 7) with a tensor product of a thin plate regression spline based on geographic coordinates of sites (longitude *lo* and latitude *la*) (Wood 2003, 2017). As the number of associated pairs and connectance are directly affected by the number of species (see correlation between network metrics in supplementary material 4) (Blüthgen et al. 2006, Besson et al. 2019), we added the number of species as a control variable for these two metrics, to estimate if the observed change in the number of associated pairs was higher or lower than expected. Similarly, connectance constrains the possible values of modularity and degree distribution (Poisot and Gravel 2014, Delmas et al. 2019), and thus connectance was added as a control variable for estimating changes in modularity and evenness of degree distribution. As the number of monitored sites changes between years, the site identity *z* was added as a random variable.

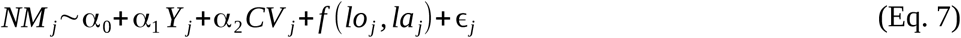

with *NM* the network metric (number of species *s*, number of associations *a*, connectance *c*, modularity *m* and evenness *e*) and *CV* the control variable (∅ for *s*, *s* for *a* and *c*, *c* for *m* and *e*) and *ε_j_∼N(0,σ)*.

## 3. Results

### 3.1 Species associations

Among the 292,796 combinations between 5,886 pairs of bird species, from four biogeographic regions and 19 habitats, we found 89,270 potential species associations (Fig. 1a). Among them, 32,491 combinations enable to reconstruct time-series with more than 500 time steps and we found 585 combinations with significantly associated species pairs (Fig. 1b). Among the 5,886 species pairs, this represents 396 (7%) associated species pairs (see association matrix in supplementary material 5), some species pairs being significantly associated in several habitats and biogeographic regions.

We found negative relationships between the propensity of a species pair to be associated and functional (−0.22 ± 0.10, Fig. 2a) and niche overlap distances (−1.73 ± 0.17, Fig. 2a). In other words, species that are more functionally similar and with higher niche overlap are more likely associated. However, we found only weak evidence of a relationships between the propensity of a species pair to be associated and phylogenetic distance (−0.09 ± 0.10, Fig. 2a and model validation in supplementary material 4) as part of the variation might be already explained by functional distance (slightly correlated with phylogenetic distance, see supplementary material 3). Concerning specific expected interactions, we found a positive relationship between the propensity of a species pair to be associated and diet during breeding season (0.23 ± 0.20, Fig. 2b), a positive relationship with similar preferred habitat (1.05 ± 0.19, Fig. 2b) and no evidence of a relationship with similar nest type (0.10 ± 0.18, Fig. 2b).

**Figure 2:**
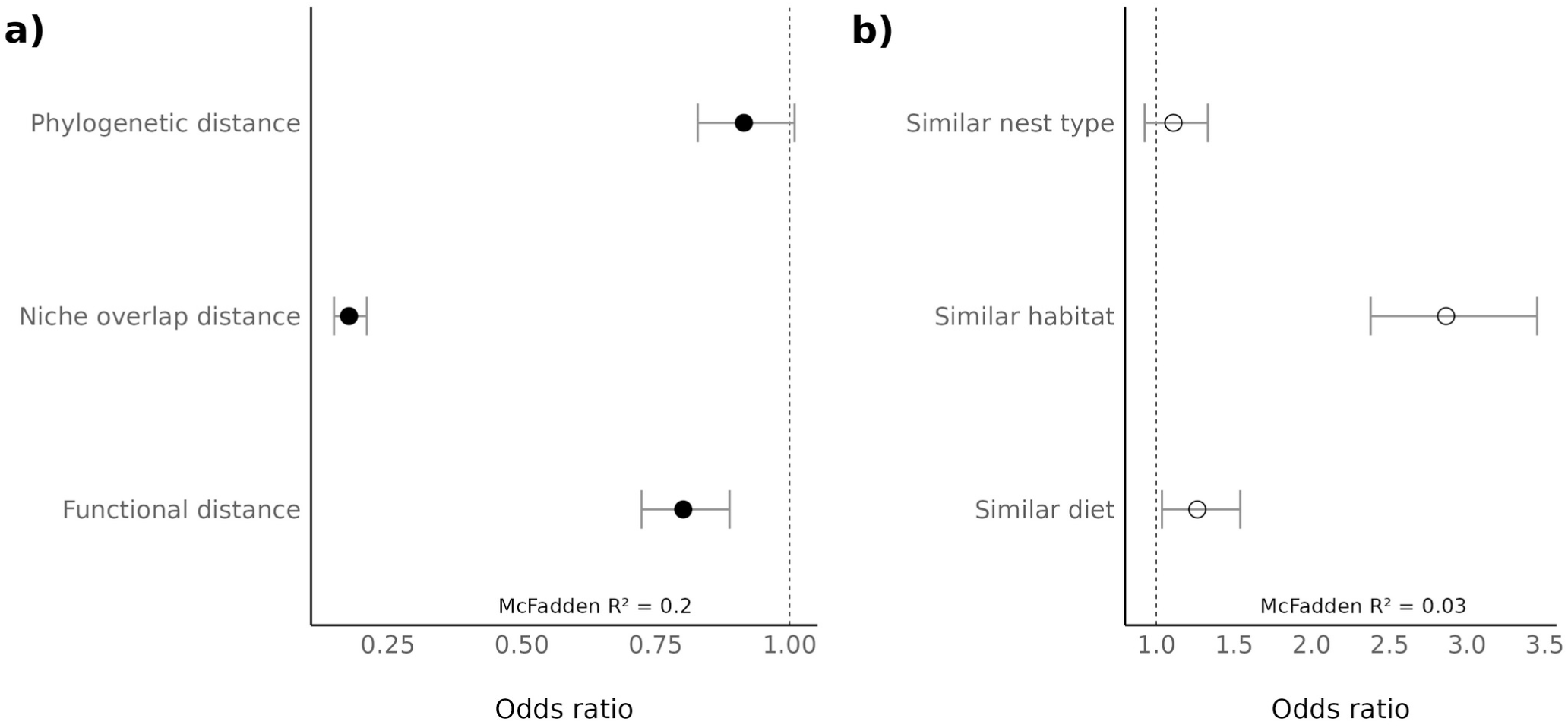
Odd ratios of GLM between propensity of a species pair to be associated and a) in black, the ecological distances (phylogenetic, niche overlap and functional distances), b) in white, specific expected interactions (similar nest, similar preferred habitat and similar diet in breeding season). 95% CI are displayed with grey bars. McFadden R² corresponds to the pseudo-R² for logistic regression models (McFadden 1974).

### 3.2 Temporal change in the interaction milieu

Breeding bird communities tend to decrease in size (*s.year^−1^*= −0.15 ± 0.02, Fig. 3a, see annual variations in supplementary material 4), increase in connectance (*c.year^−1^* = 5.6.10^−4^ ± 2.3.10^−4^, Fig. 3c) and decrease in evenness of the degree distribution (*e.year^−1^* = −1.1.10^−4^ ± 0.5.10^−4^, Fig. 3d) over the studied period (2001-2017). This is verified for farmland and forest communities that represent the majority of communities (55.3% in farmland, 29.7% in forest, 7.9% in natural open land and 7.1% in urban areas). For farmland and forest communities, the increase in connectance is related to the decline in the number of species, whereas the number of associations relatively increases compared to the number of species in natural open land communities. For the interaction milieu in urban communities, the only significant change is the decline of the number of species (Fig. 3a), the other metrics staying constant.

**Figure 3:**
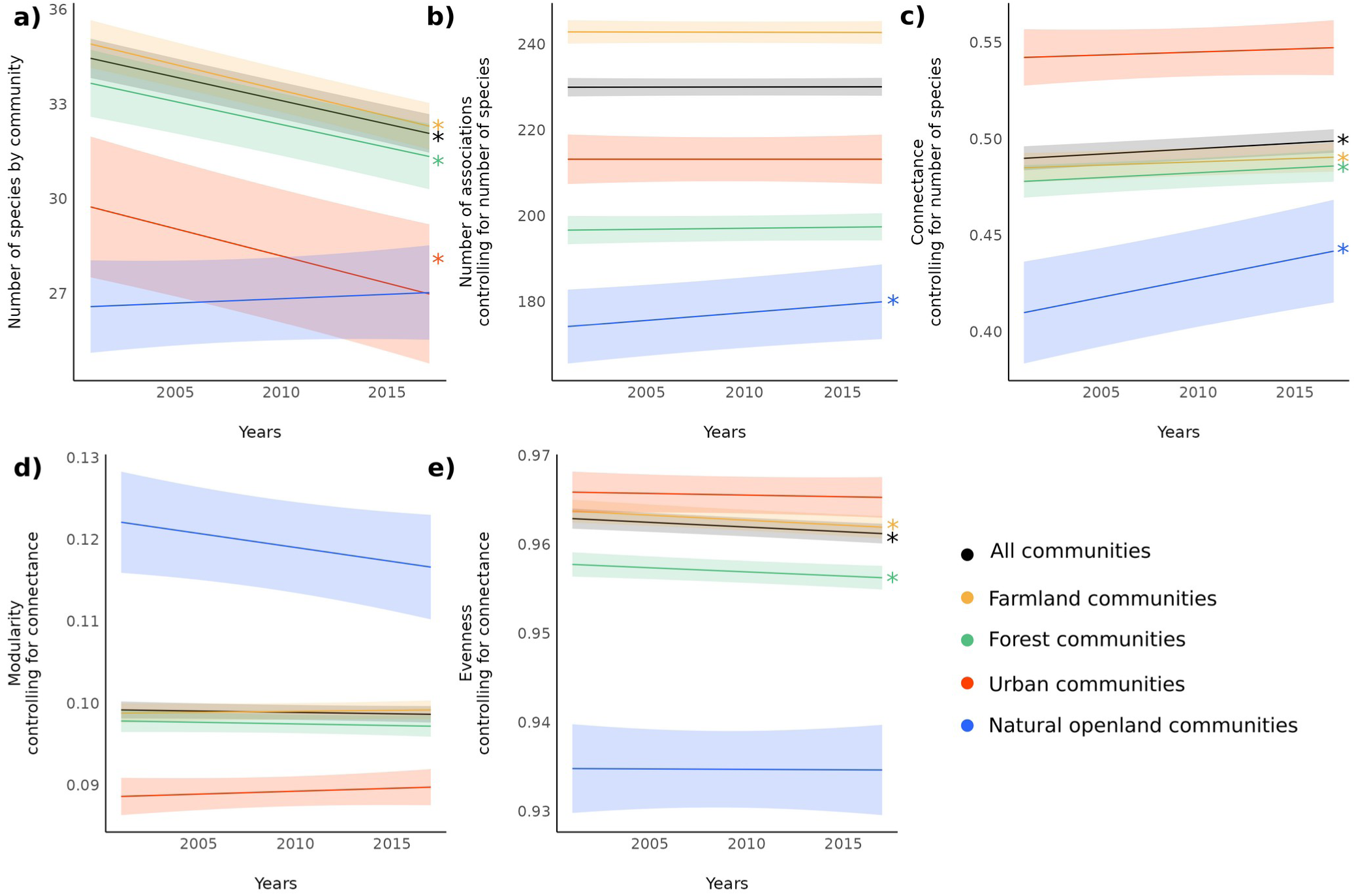
Temporal changes in the interaction milieu of bird communities between 2001 and 2017. a) Temporal change in the number of species *s*, b) temporal change in the number of associations *a* (while controlling for the number of species), c) temporal change in connectance *c* (while controlling for the number of species), d) temporal change in modularity *m* (while controlling for connectance), e) temporal change in evenness of the degree distribution *e* (while controlling for connectance). Black dot is for all communities, colours for communities from each habitat: yellow for farmland, green for forest, red for urban, blue for natural open land. 95% CIs are displayed and significant changes are shown by ⁎.

Consequently, the interaction milieu from the start of the period (Fig. 4a) is modified by a change in both the number of species (Fig. 4b) and the topology of relationships between species (Fig. 4c). At the level of the community, the density of associated pairs (i.e. connectance, Fig. 3c) is higher, which is mainly due to the reduction in the number of species (Fig. 3a), not to a relative increase of the number of associated pairs (Fig. 3b). The intensity of division of the community into sub-groups (i.e. modularity) remains stable when accounting for the density of associations. Thus, the major change is at the species level, where the probability distribution of the number of association by species (i.e. evenness) tends to become more skewed (i.e. shifting from species with a similar number of association to species with few or multiple associations). The trend over the period studied is from large communities with evenly distributed associations (Fig. 4a) to smaller communities with a similar relative number of associations but unevenly distributed (Fig. 4d), which an focus on species only would not have revealed.

**Figure 4:**
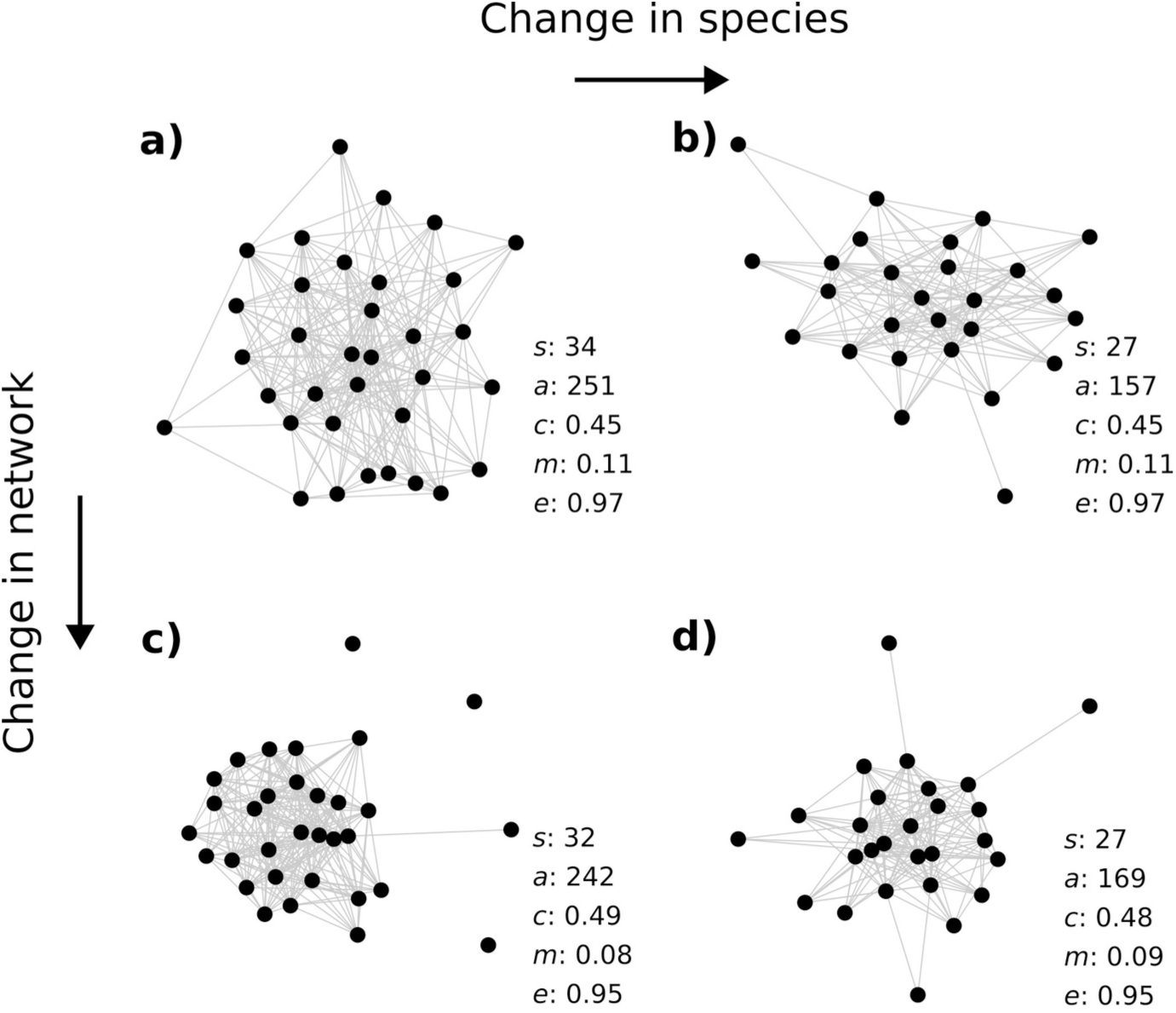
Observed examples of bird communities illustrating the topological changes in the interaction milieu between 2001 and 2017 with the values for each network metric. From a) a community with a given number of species *s* and network metrics (number of associations *a*, connectance *c*, modularity *m* and evenness *e*), b) network metrics may not be affected by a change in the number of species, c) conversely, a stable number of species may hide a change in network structure. d) On average, the observed changes between 2001 and 2017 in bird communities show an interaction milieu with a lower number of species and a similar relative number of associations that are becoming less evenly distributed, as illustrated by the observed network displayed.

## 4. Discussion

### 4.1 Ecological relevance of associated species pairs

The species pairs we have identified as associated share similar functional characteristics or habitat requirements. The significant relationships with functional and niche overlap distances were in line with our hypothesis based on trait-based assembly theory and empirical results on birds (Mönkkönen et al. 2017) and suggest that our approach produced ecologically meaningful associated species pairs. The absence of a significant relationship between associated species pairs and phylogenetic distance can be linked to the phylogenetic diversity already encompassed in functional distance (see supplementary material 3). Despite this result and the strong filters applied in the estimation of associated pairs, some of the associated pairs can still be due to a common response to an environmental variable in space and time (Pollock et al. 2014) and thus an absence of association does not imply an absence of interaction between two species (Cazelles et al. 2016). Yet, we provide an in-depth validation of the overall set of associated pairs using the ecological distances and confirm this by looking at expected interacted pairs of species based on specific interactions during the breeding season (corresponding to the period of the survey) for food and nest location. The similarity in nest location is not significant, which may be due to the fact that competition would be the main expected interaction from this shared characteristic (Martin et al. 2004). On the contrary, a similar diet, that significantly increases the propensity to be associated, could induce both repulsion and attraction (Seppänen et al. 2007, Gil et al. 2017) and thus is expected to induce interactions in a wider range of species pairs and to be more easily detected. This result supports the findings based on functional and niche overlap distances (and see supplementary material 4 for additional detail on specific associations).

### 4.2 Specificity of the interaction milieu

The changes in the interaction milieu of bird communities estimated from the associated species pairs show a decrease in the number of species and the evenness of the number of associations by species as well as an increase in the density of association, with urban and natural open land communities that differ slightly from this general pattern. These changes in the biotic background of interaction in bird communities remain estimated from species links that are coarse and static, whereas ecological interactions are known to be changing though time (Ramos-Robles et al. 2016, Suzuki et al. 2023), due in particular to changes in environmental conditions (Lurgi et al. 2012, Poisot et al. 2015). In addition, the interpretation of changes in the interaction milieu is limited by the absence of formal identification of the interaction nature, compared to explicit ecological networks (e.g. mutualistic plant-pollinators, food webs). Our finding can therefore only be compared by analogy to changes observed in ecological networks with specific interactions such as plant-pollinator networks and food webs. In particular, the changes observed in just two decades in the interaction milieu of bird communities can be considered as marked compared to analogous communities in which frugivore-plant, pollination networks and food webs are reported to be stable (Petanidou et al. 2008, Kaartinen and Roslin 2012, Plein et al. 2013). Plant-pollinator networks, food webs and research on bird flocks (closely related to intra-guild networks) may however bring clues to interpret the observed changes in the interaction milieu of bird communities.

### 4.3 Changes in the interaction milieu and consequences for conservation

Based on an analogy between these ecological networks and bird communities (Lane et al. 2014, Mokross et al. 2014), we can assume in particular that specialists species are supposed to be involved in fewer but stronger interactions (Rigal et al. 2022). If changes in connectance alone are not informative for conservation (Heleno et al. 2012), the increase in connectance is related here to a reduction in the number of species in bird communities that could be interpreted in line with the observed decline of habitat specialist species (Devictor et al. 2008b). Higher connectance can improve network persistence facing perturbation (Besson et al. 2019), which might be the case if all the fragile species and interactions are already wiped out while generalists involved in more associations relatively expand (Godet et al. 2015). However, if this connectance increase is at the cost of species removal, according to the biodiversity-ecosystem functioning theory (which is relevant inside a guild), this will result in a declining efficiency of resource use (Loreau 2010). Based on our results, such a decline in community functioning is therefore expected for forest and farmland communities, in which the change in connectance is explained by the decline in species number. Conversely, the structure and connectance of communities in natural open land are principally modified by the replacement of less associated species by more associated ones, expected to be generalists.

An increase in connectance also induces a decline in modularity (Delmas et al. 2019) which has been the case in bird communities, although modularity change is not stronger or weaker than expected (i.e. the modularity does not change through time when we control for changes in connectance Fig. 3d). A modular structure is known to reduce the impact and propagation of a perturbation throughout the network (Stouffer and Bascompte 2010). Therefore, considering consequences for the interaction milieu, a decline in modularity might affect community ability to cope with environmental changes and induce a higher sensitivity to cascade and secondary extinctions.

The observed decline in evenness of association degree distribution can be more directly interpreted. Low evenness in the distribution of interaction strength can be often observed in ecological networks, due to the numerous weak interactions (Bascompte et al. 2005) known to stabilise community dynamics. However, when focusing on the presence, not the strength, of a link, the evenness of the distribution is higher in preserved or restored communities than in degraded habitats (Tylianakis et al. 2010, Fisogni et al. 2021), not only in food webs but also in avian flocks (Mokross et al. 2014). It is therefore likely that the observed decline in evenness, which is higher than expected by the observed change in connectance, is the footprint of habitat degradation experienced by bird communities. The observed decline in evenness in forest and farmland communities may therefore be compared with the well-studied decline in species dynamics and changes in community composition over the past decades in these environments (Brotons and Jiguet 2010, Gaüzère et al. 2020, Rigal et al. 2023).

Although our study focuses on the interaction milieu of bird communities, this should not understate the importance of the marked decline observed in the size of these communities. Beyond the species loss that is an issue per se, a decline in species diversity and abundance is expected to affect interaction frequency as the encounter rates become lower. This could lead to the loss of some interactions while the two species are still present in the vicinity (Staniczenko et al. 2017, Tylianakis and Morris 2017). Yet, species remains a central unit for conservation purposes and our results support the fact that change in ecological network structure should be analysed in regard with change in species number and identity (Heleno et al. 2012).

Changes in the interaction milieu of bird communities highlighted in this study are crucial to understand how communities will be able to respond to future perturbations, in particular if community functioning is affected by changes in the network layout (Besson et al. 2019). Community responses are affected by the loss of species (Loreau and De Mazancourt 2013) which is, additionally, deepened by the rewiring (Tylianakis and Morris 2017) or the vanishing of associations (Gilljam et al. 2015). These rapid modifications of bird biodiversity, that would have been hidden by a species-centred approach, are expected to affect the stability of communities and their capacity to cope with rapid and multiple global changes.

## Data availability

All the analyses are conducted using the R software v.4.2.2 (R Core Team 2018). Code and data are available on Zenodo https://zenodo.org/doi/10.5281/zenodo.12784005 (Rigal et al. 2024).

## Author Contributions

S. Rigal, V. Devictor and V. Dakos conceived the ideas and designed methodology; S. Rigal analysed the data and led the writing of the manuscript. All authors contributed critically to the drafts and gave final approval for publication.

