## supplementary material for "Tracking changes in birds’ interaction milieu"

**Supplementary material 1: Habitat class and potential associated pairs**

| **Merged classes** | **Habitat classes** |
| --- | --- |
| Forest | Deciduous woodland |
|  | Coniferous woodland |
|  | Mixed woodland |
| Natural open land | Young forest |
|  | Heath |
|  | Coppice |
|  | Dry natural meadow |
|  | Moorland |
|  | Marshland |
|  | Near open water |
|  | Bare rocks |
| Farmland | Ploughed meadow |
|  | Unploughed meadow |
|  | Mixed farmland |
|  | Open-field |
|  | Permanent crop |
| Urban | Urban settlement |
|  | Suburban settlement |
|  | Rural settlement |

Table S1: Habitat classes summarized from FBBS habitat types.

We estimated potential associated pairs of species from bird co-abundance data (Morueta-Holme et al. 2016) for each of the four biogeographic regions and for each of the 19 habitats using the four following steps (Fig. 3).

Step 1. In order to limit the influence of phylogeography and habitat features on species associations, we first grouped the data by biogeographic region (Continental, Atlantic, Mediterranean, Alpine) and by habitat (19 habitat classes, Table S1).

Step 2. In each biogeographic region and habitat, we used the log-transformed co-abundance data (to obtain normally distributed data) to calculate observed associations as partial correlations between each pair of species (Schäfer and Strimmer 2005) as follows (Eq. 1):

$\text{Pc}\left( O \right)_{\text{i,j}}=\frac{-\sum(O)}{\sqrt{\sum(O)\text{.}\sum(O)}}$	(1)

with *O* the matrix of observed abundance (species x sites), *Pc(O)_i,j_* the partial correlation between species *i* and *j*, and *Σ_i,j_^-1^* the value for species *i* and *j* of the inverse of the covariance matrix. This approach partially removes the indirect effects of other co-occurring species on the estimated association between the two considered species by focusing on the conditional association (Harris 2016).

Step 3. Partial correlations can be affected by species commonness, since common species have higher probabilities to co-occur than less abundant species because of a higher representativeness in the data (Blüthgen et al. 2008). To correct this bias, we computed partial correlations on 1000 random co-abundance datasets obtained by keeping constant the total number of individuals in a given sampling point, and assuming that the probability for a species to occur in a given sampling point was proportional to its frequency in the dataset. We then calculated standardised effect sizes of partial correlations between species *i* and *j* (*SES_i,j_)* as follows (Eq. 2):

$\text{SES}_{\text{i,j}}=\frac{\text{Pc}\left( O \right)_{\text{i,j}}-\mu\left( \text{Pc}\left( N \right)_{\text{i,j}} \right)}{\sigma\left( \text{Pc}\left( N \right)_{\text{i,j}} \right)}$	(2)

where *Pc(O)_i,j_* is the observed partial correlation between species *i* and species *j*, *μ(Pc(N))_i,j_* and *σ(Pc(N))_i,j_* the mean and standard deviation of partial correlations from the 1000 randomly sampled datasets.

Step 4. In order to identify *significant* associations, we calculated a two tail p-value for each pairwise association using the rank of the observed association in the Gaussian distribution of null associations obtained from step 3. That is, we determined the number of replicates for which the absolute value of the observed partial correlation is greater than the absolute null partial correlation (p-values were corrected for multiple comparisons following Benjamini and Hochberg (1995)). Potential associated species pairs therefore corresponded to species pairs *i,j* with a *SES_i,j_* for which the adjusted p-values was below 0.05.

**Supplementary material 2: CCM calculation details**

We determined the existence of the interaction as a causal relation between species time-series estimated by Convergent Cross-Mapping (CCM) (Sugihara et al., 2012). The idea behind this approach is that if two species are causally linked, their time-series have a common attractor manifold. By reconstructing the time-lag manifolds of X and Y, respectively M_X_ and M_Y_, it is possible to test whether M_X_ and M_Y_ are empirical reconstructions of the same manifold M. This is done by testing if, for the nearest neighbours of a point in M_Y_, their corresponding points (identified through the time index) are also nearest neighbours of the corresponding point in M_X_ (Fig. S1A-B). That is, one time-series can estimate the state of the other. By testing how well values of one time-series can predict the state of the other, CCM allows to retrieve the causality between species time-series (Fig. S1C). However, this method requires time-series with at least 30 records. Thus, we used the multispatial CCM method (R package multispatialCCM) (Clark et al., 2015), allowing to determine causality in short times-series. This method uses information given by spatial replicates of short time-series to apply CCM on long enough pseudo-time-series.

CCM requires a three step approach. 1) According to Taken's theorem (Takens, 1981), the core of the CCM method, it is possible to reconstruct *M_X_* using several time-lags of the time-series of X. Hence the first of the three steps is to find the best embedding dimension *E* to precisely map the original manifold *M*. As time-series used in multispatial CCM come from multiple sites, *E* can not be higher than the minimum number of time steps by site *n* (*E* ⩽ *n-1* as one time step must be kept for prediction, see step 2). This is why we used time-series with at least six consecutive time steps, allowing *E* to be less than or equal to 5. 2) Once the best *E* is determined, one needs to check whether dynamics are strongly influenced by noise, leading to a purely random system, or not. To do so, a part of the observations are used to make predictions for future and increasingly far observations and their predictive power is estimated. If the system is non-linear and not driven by an important stochastic noise, the predictive power should decrease with temporal distance. 3) Once E is determined and non-linearity verified, CCM can be applied.

We applied multi-scale CCM with 300 replicates for the bootstrap routine due to computational time. This gave us species pairs for which the time-series of one was causally related to the time-series of the other. We considered there was no temporal association between two species if none was causally related to the other, and a temporal association between two species if one of the species was causally related to the other or if each of the two species were causally related to the other. We considered the species composing such pairs as being associated.

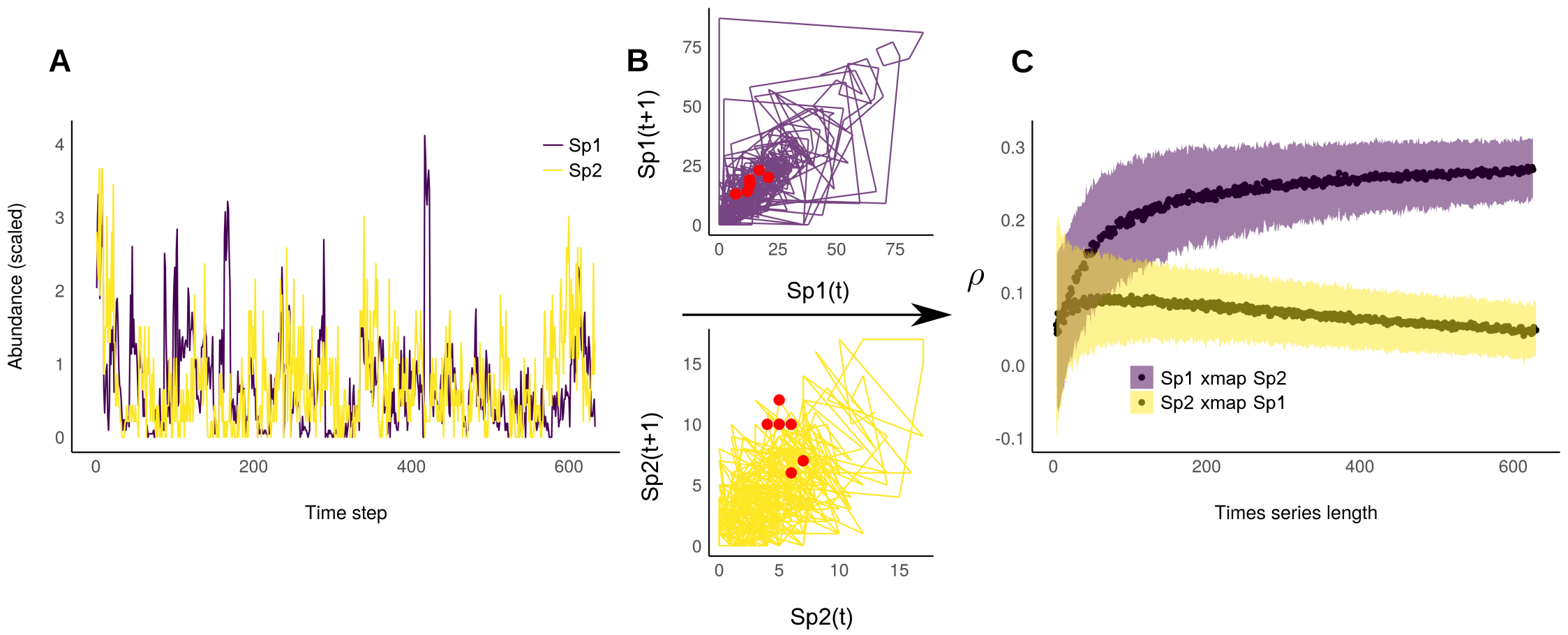
Figure S2: Determining and quantifying species temporal associations. A) Species time-series reconstructed from several locations (*Sp1: species 1, Sp2: species 2*). B) Attractor manifold of *Sp1* and *Sp2*. As an example, red dots correspond to attractor states for the same six temporal steps. They are grouped in manifolds of *Sp1* and *Sp2*, therefore *Sp1* and *Sp2* probably shared a common manifold and this is evaluated using Convergent Cross Mapping (CCM). C) Cross Map Skill *ρ* of *Sp1* and *Sp2* with increasing time-series length. This allows to find out the significance of the temporal association between *Sp1* and *Sp2*.

**Supplementary material 3: Functional, phylogenetic and niche overlap distances**

1. Niche overlap distance

Figure S3A: Niche overlap (defined as the inverse of niche overlap distance and scale between 0 and 1 for graphical purposes) between species. Dark blue corresponds to high habitat overlap between species and ligh
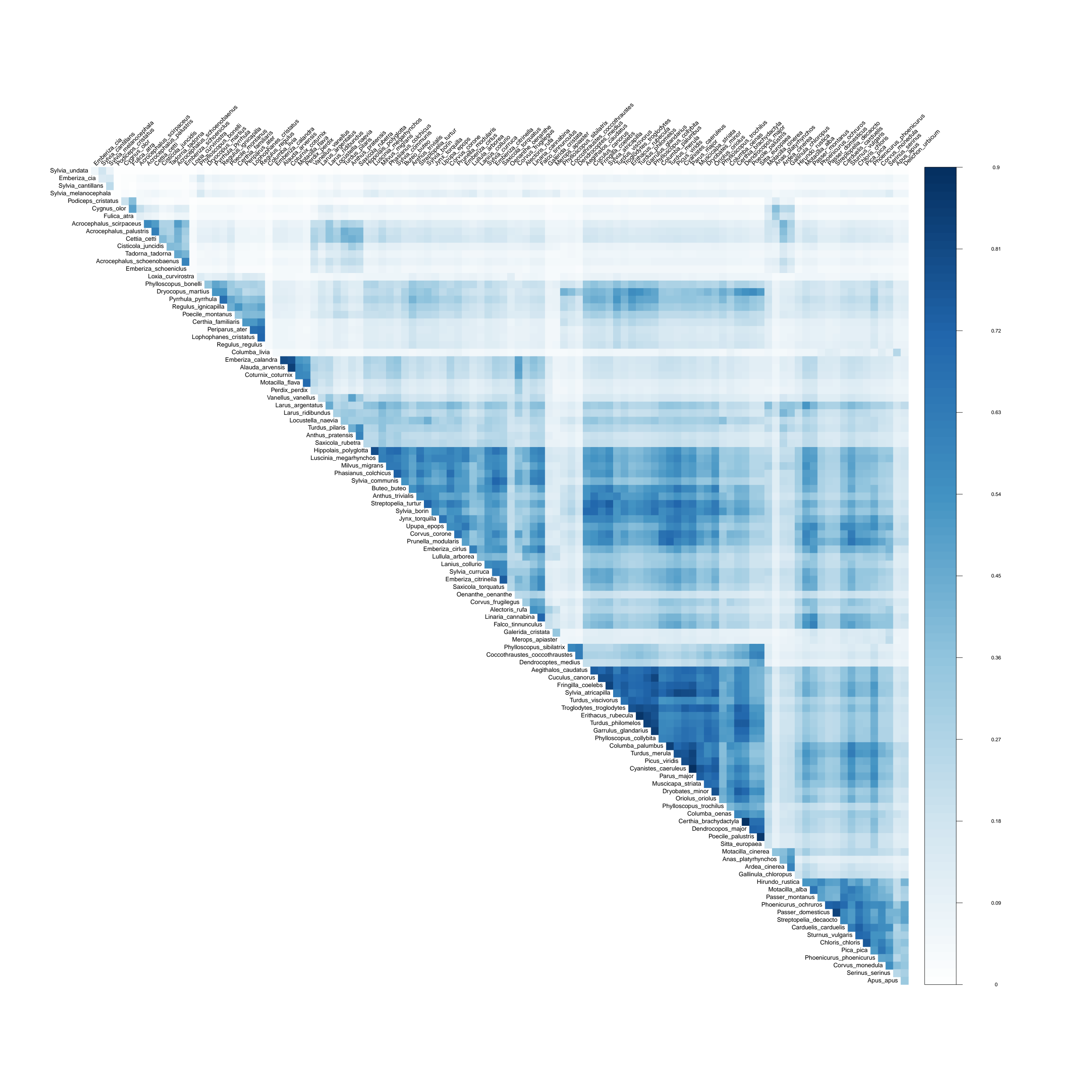
t blue to no habitat overlap between species. Species are regrouped using hierarchical clustering order. From left to right and up to down, the first triangle regroups marsh species (*Acrocephalus scirpaceus, Acrocephalus palustris, Cettia cetti, Cisticola juncidis, Tadorna tadorna, Acrocephalus schoenobaenus, Emberiza schoeniclus*), the second regroups woodland specialist species (*Loxia curvirostra, Phylloscopus bonelli, Dryocopus martius, Pyrrhula pyrrhula, Regulus ignicapilla, Poecile montanus, Certhia familiaris, Periparus ater, Lophophanes cristatus, Regulus regulus*), then farmland species (*Emberiza calandra, Alauda arvensis, Coturnix coturnix, Motacilla flava, Perdix perdix*), then species from open areas (*Hippolais polyglotta, Luscinia megarhynchos, Milvus migrans, Phasianus colchicus, Sylvia communis, Buteo buteo, Anthus trivialis, Streptopelia turtur, Sylvia borin, Jynx torquilla, Upupa epops, Corvus corone, Prunella modularis, Emberiza cirlus, Lullula arborea, Lanius collurio, Sylvia curruca, Emberiza citrinella, Saxicola torquatus, Oenanthe oenanthe, Corvus frugilegus, Alectoris rufa, Linaria cannabina, Falco tinnunculus*), then woodland generalist species (*Phylloscopus sibilatrix, Coccothraustes coccothraustes, Dendrocoptes medius, Aegithalos caudatus, Cuculus canorus, Fringilla coelebs, Sylvia atricapilla, Turdus viscivorus, Troglodytes troglodytes, Erithacus rubecula, Turdus philomelos, Garrulus glandarius, Phylloscopus collybita, Columba palumbus, Turdus merula, Picus viridis, Cyanistes caeruleus, Parus major, Muscicapa striata, Dryobates minor, Oriolus oriolus, Phylloscopus trochilus, Columba oenas, Certhia brachydactyla, Dendrocopos major, Poecile palustris, Sitta europaea*), waterfowl (*Motacilla cinerea, Anas platyrhynchos, Ardea cinerea, Gallinula chloropus*) and finally periurban species (*Hirundo rustica, Motacilla alba, Passer montanus, Phoenicurus ochruros, Passer domesticus, Streptopelia decaocto, Carduelis carduelis, Sturnus vulgaris, Chloris chloris, Pica pica, Phoenicurus phoenicurus, Corvus monedula, Serinus serinus, Apus apus*).

2. Relationship between niche distances

Functional, phylogenetic and niche overlap distance are not totally independent as they were partly built on similar information as shown by the relationships between those three distances (Fig. S3C). However, they were also complementary indices. Functional information captured by functional distance was not completely redundant with functional information carried by phylogenetic relationships (Fig. S3B and S3C). Niche conservatism implied a part of redundancy between phylogenetic distance and functional distance but niche exclusion process made these two indices complementary. Habitat requirement leading to niche overlap, specifies one of the many aspects taken into account in functional distance. These two indices also were complementary as they captured much different variabilities. Niche overlap successfully regrouped specialists from a given habitat together and more generalist species together (Fig. S3A).

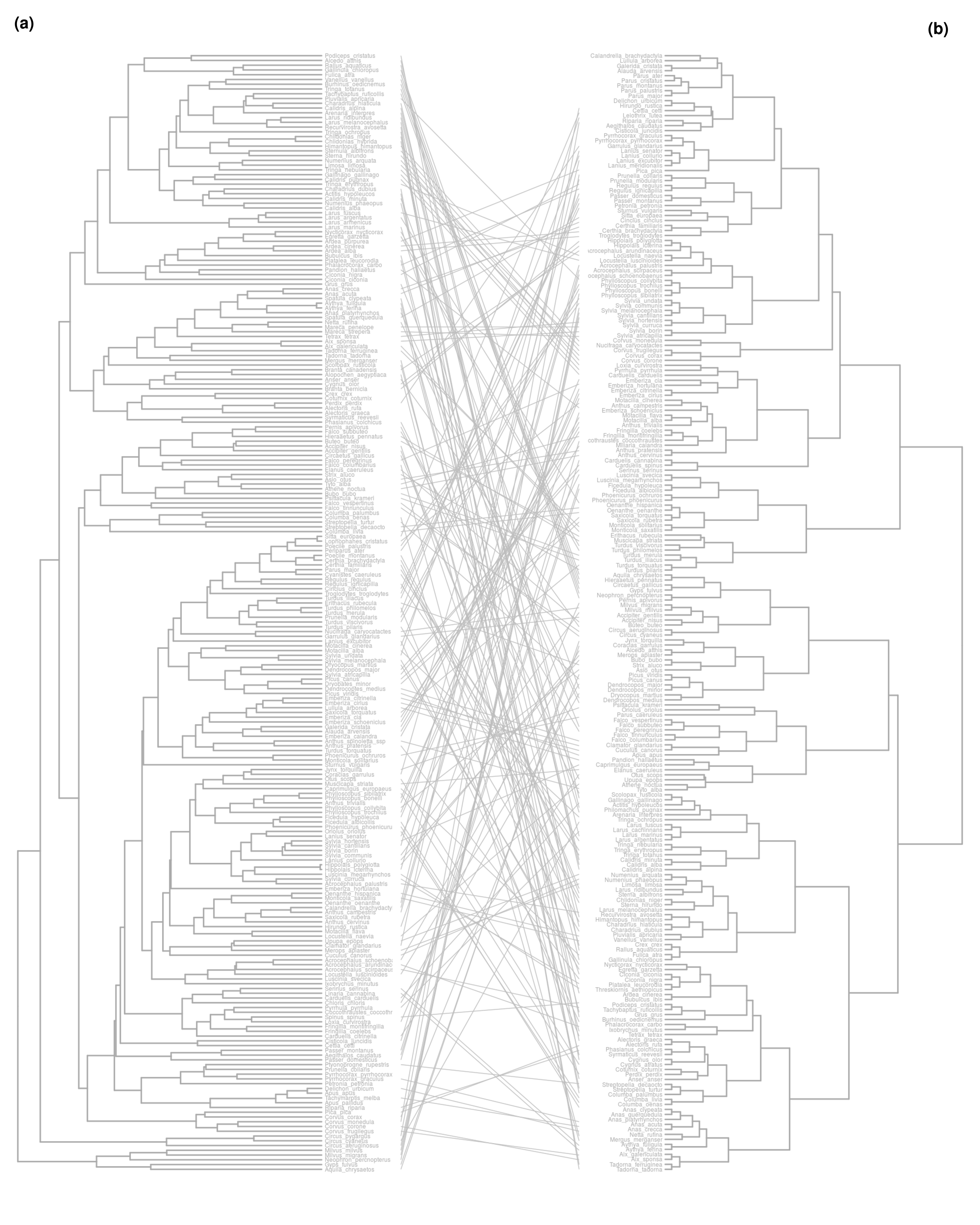
Figure S3B: Comparison between a) the functional distance tree and b) the phylogenetic distance tree.

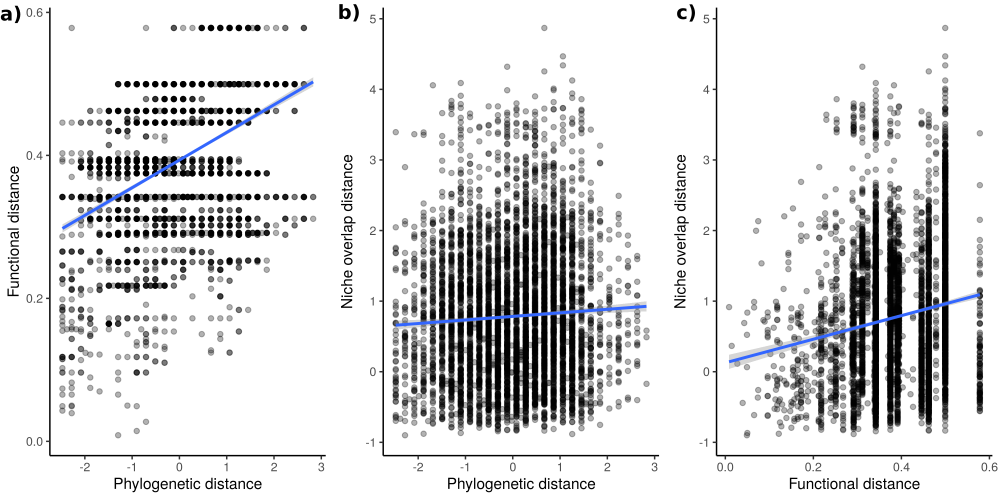

Figure S3C: Distance coplots. A) Functional versus phylogenetic distances. B) Niche overlap versus phylogenetic distance. C) Niche overlap versus functional distance.

**Supplementary material 4: Correlation between network metrics, annual variation and model validation**

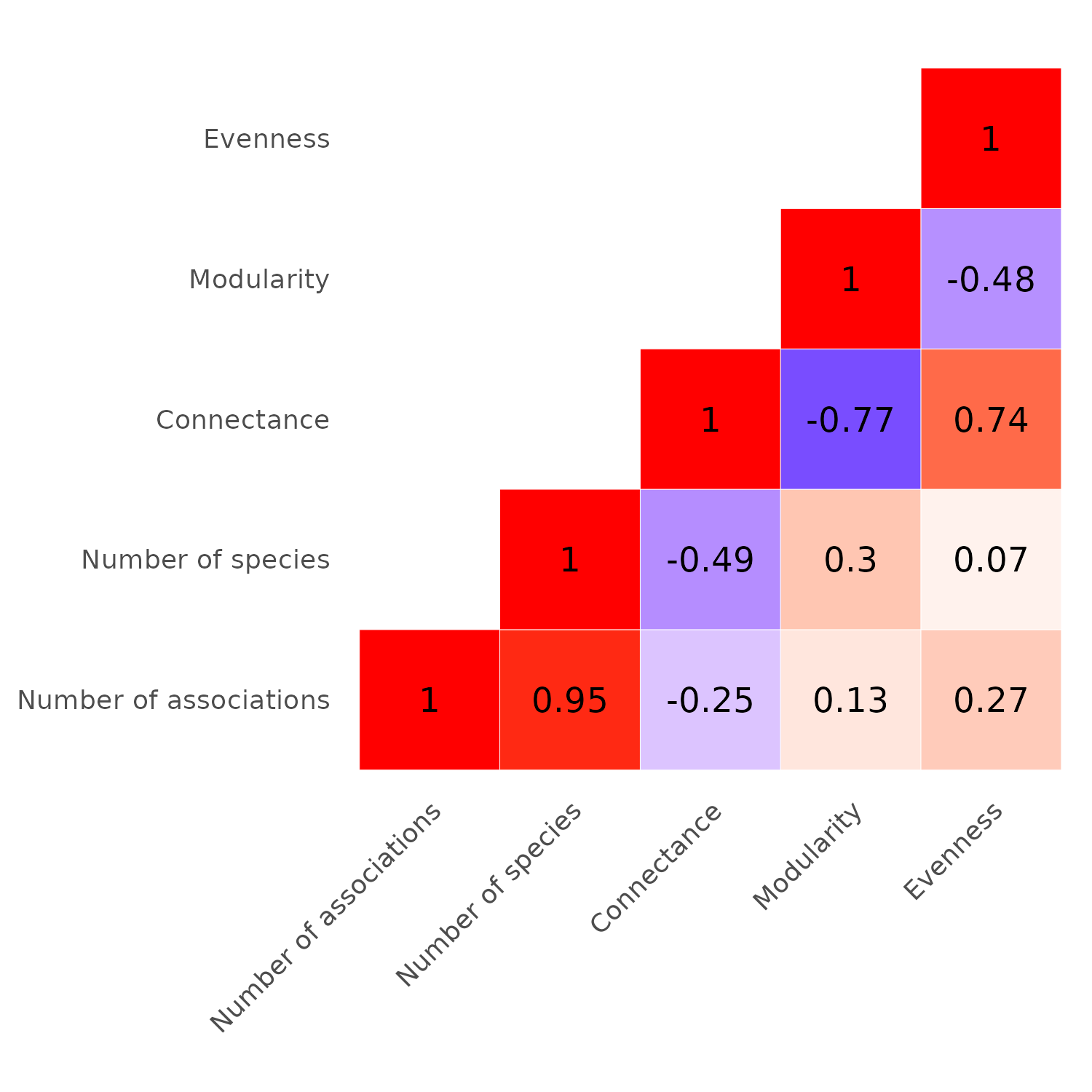
Figure S4A: Pearson correlation between number of associations, number of species, connectance, modularity and evenness in bird communities from 2001 to 2017.

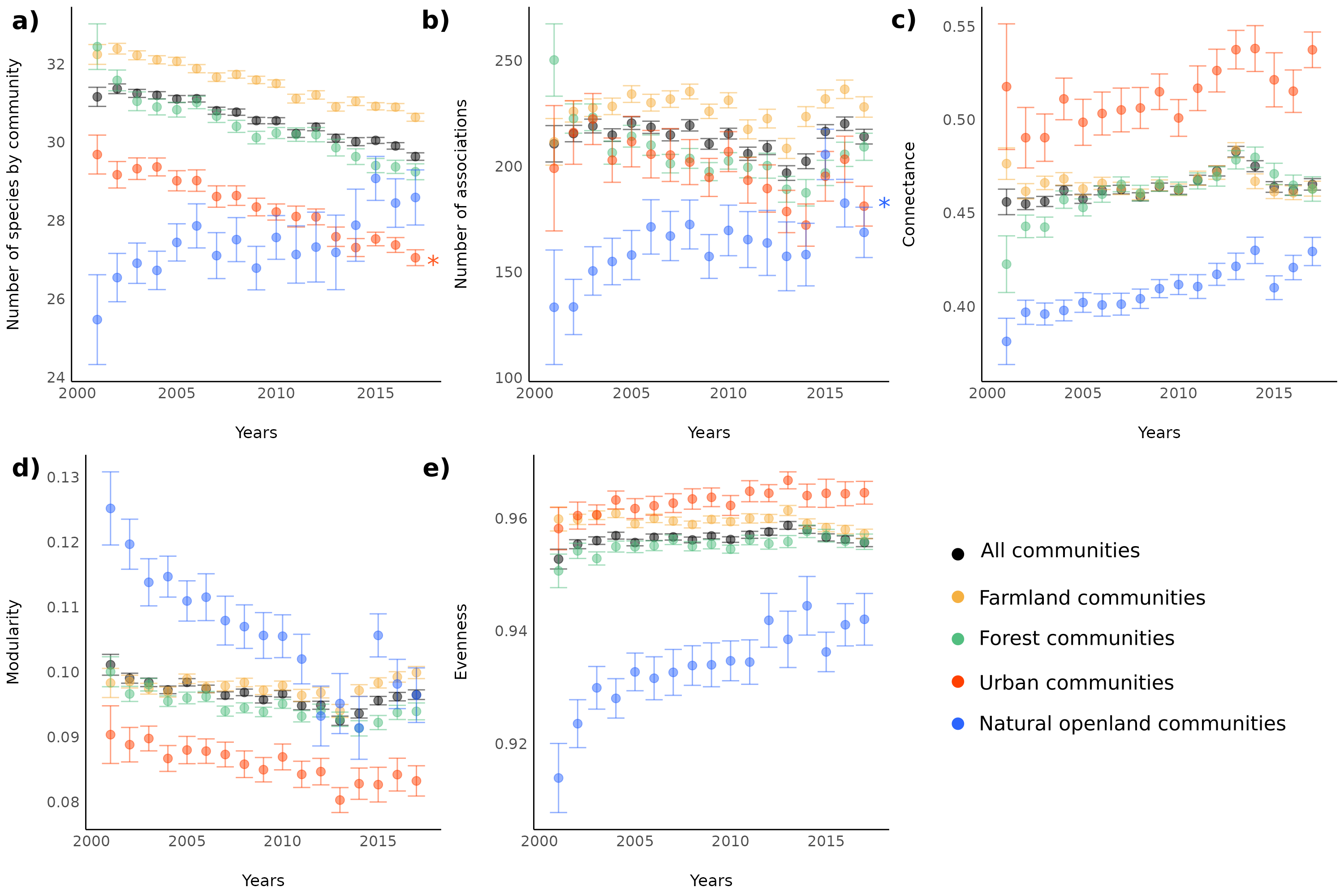
Figure S4B: Annual variation in bird communities from 2001 to 2017 in a) the number of species, b) the number of associations, c) connectance, d) modularity and e) evenness of the degree distribution. Black dot is for all communities, colours for communities from each habitat: yellow for farmland, green for forest, red for urban, blue for natural open land. 95% CIs are displayed.

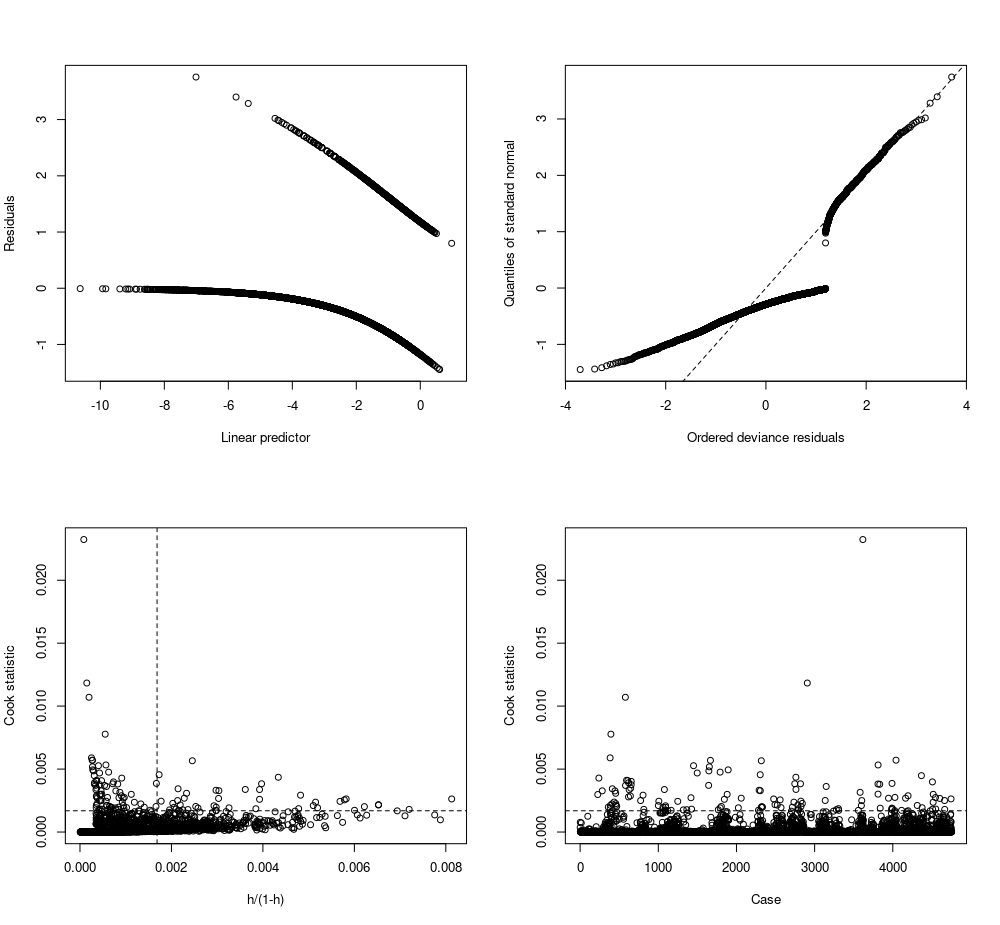
Figure S4C: Diagnostic plots for the GLM model with propensity of species association linked to functional, phylogenetic and niche overlap distances.

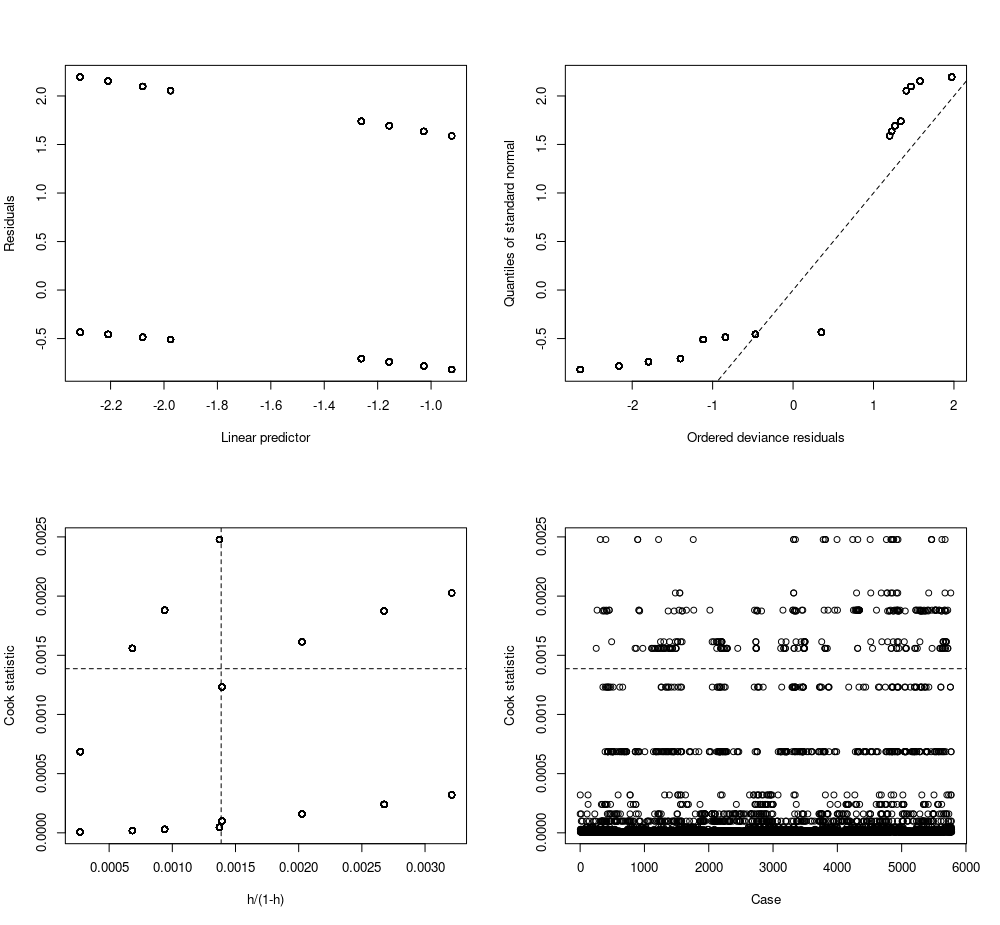

Figure S4D: Diagnostic plots for the GLM model with propensity of species association linked to similarity in nest type, diet during breeding season, and preferred habitat.

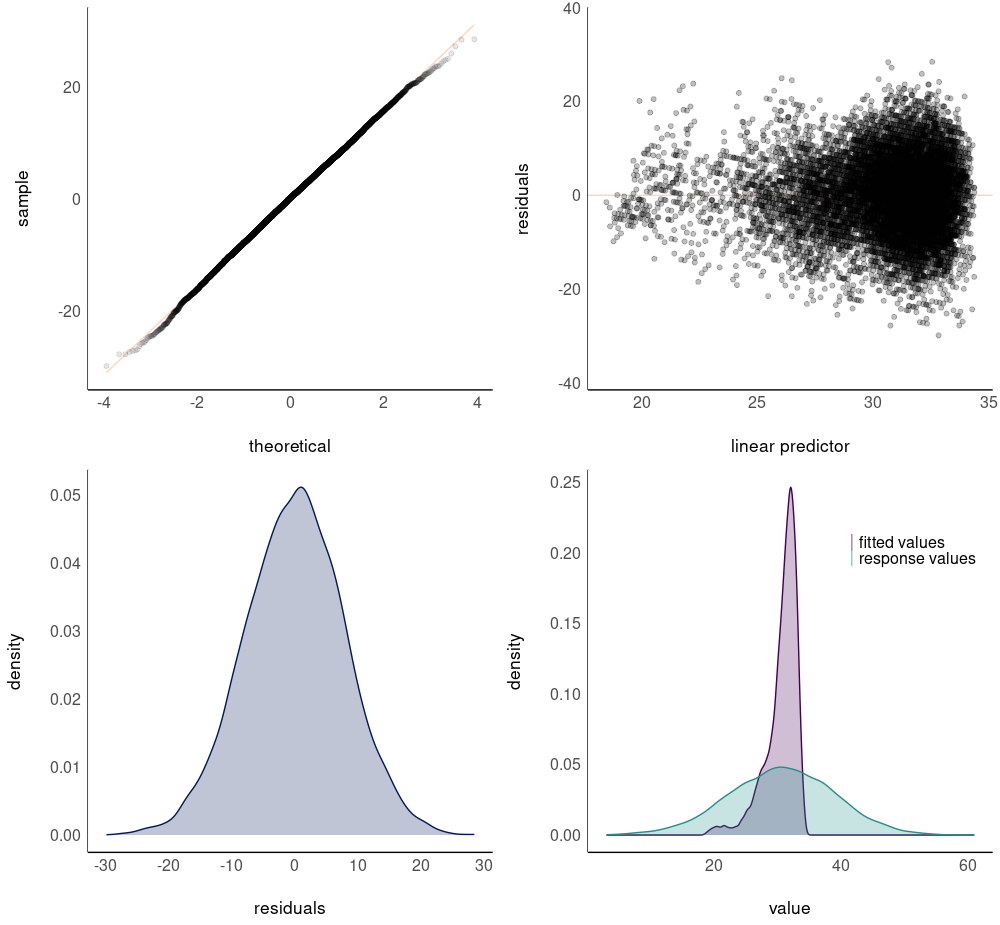
Figure S4E: Diagnostic plots for the GAMM model with network size (number of species) as response variable and years as explanatory variable and connectance as control variable.

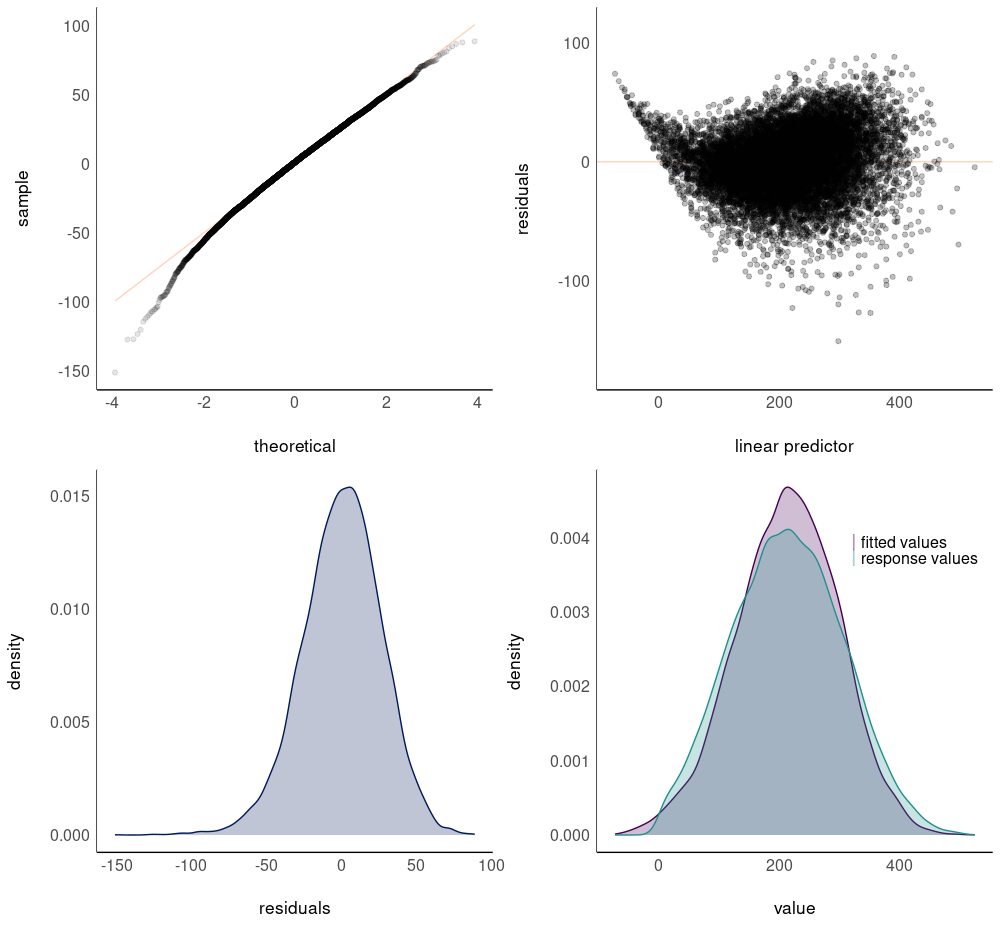
Figure S4F: Diagnostic plots for the GAMM model with number of associations as response variable and years as explanatory variable and network size as control variable.

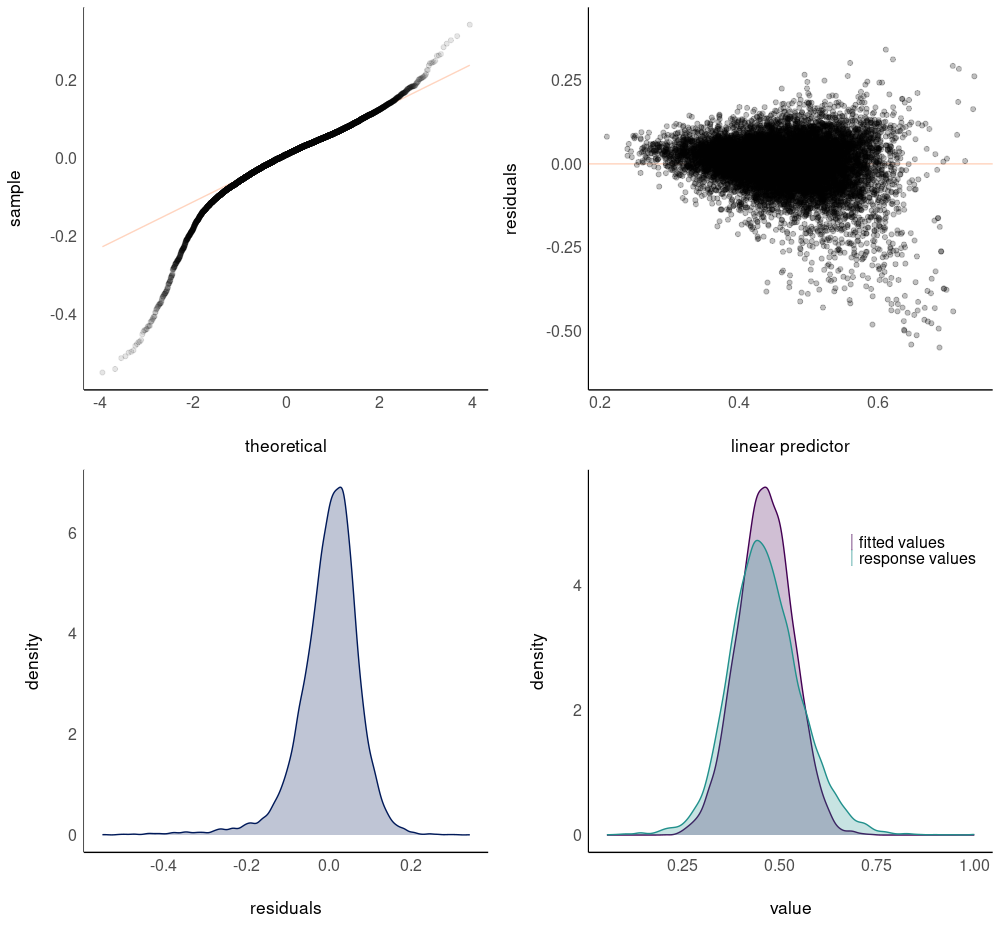
Figure S4G: Diagnostic plots for the GAMM model with connectance as response variable and years as explanatory variable and network size as control variable.

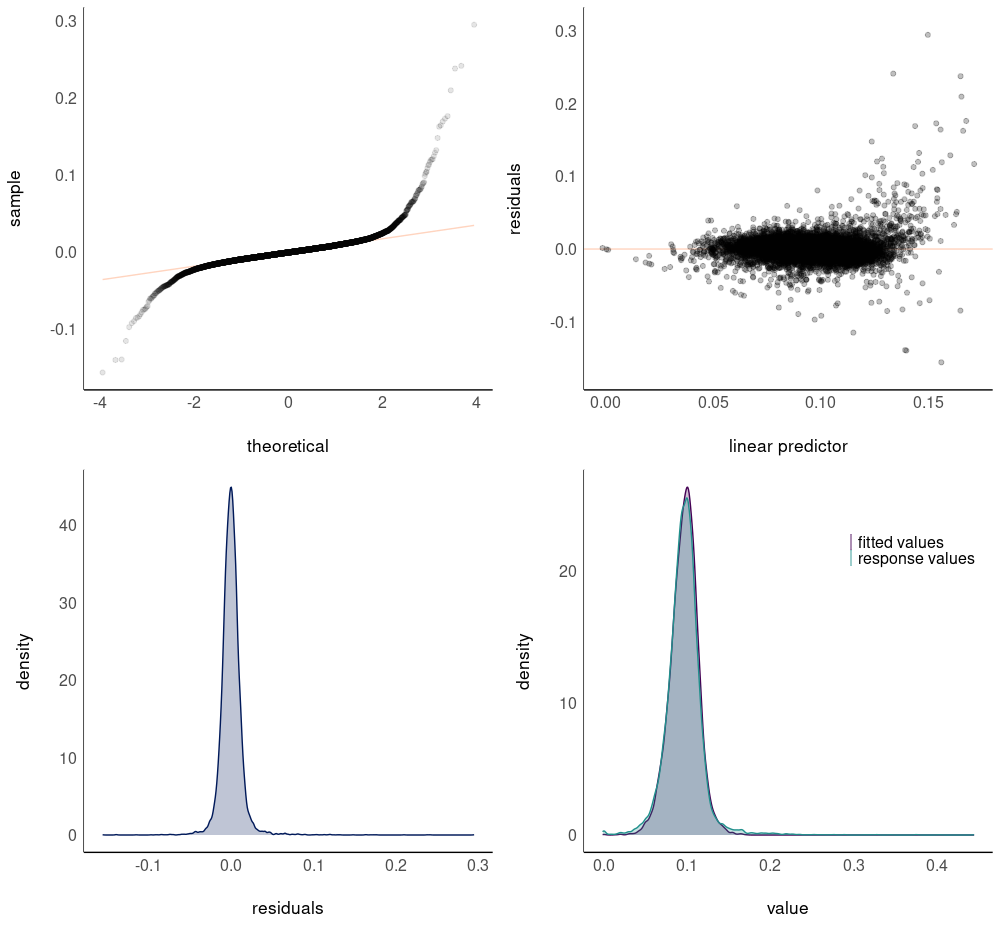
Figure S4H: Diagnostic plots for the GAMM model with modularity as response variable and years as explanatory variable and connectance as control variable.

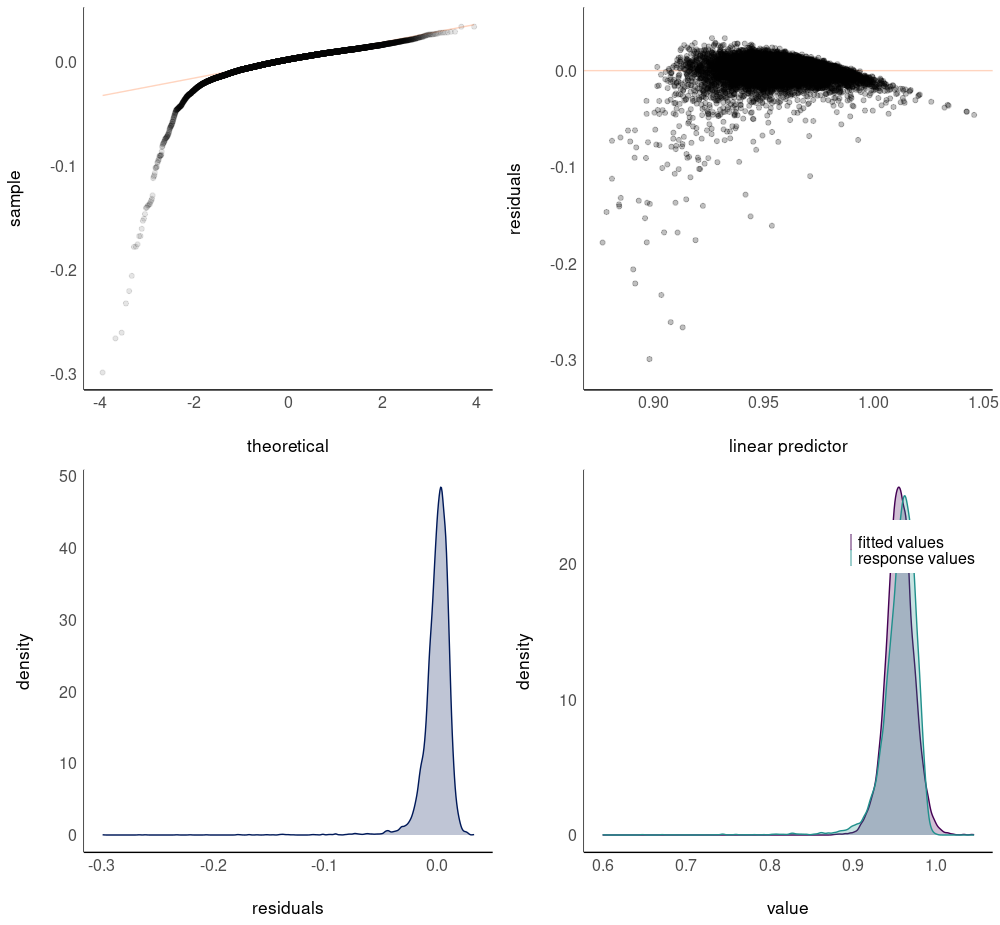

Figure S4I: Diagnostic plots for the GAMM model with evenness as response variable and years as explanatory variable and connectance as control variable.

Supplementary material 5: Details on species associations

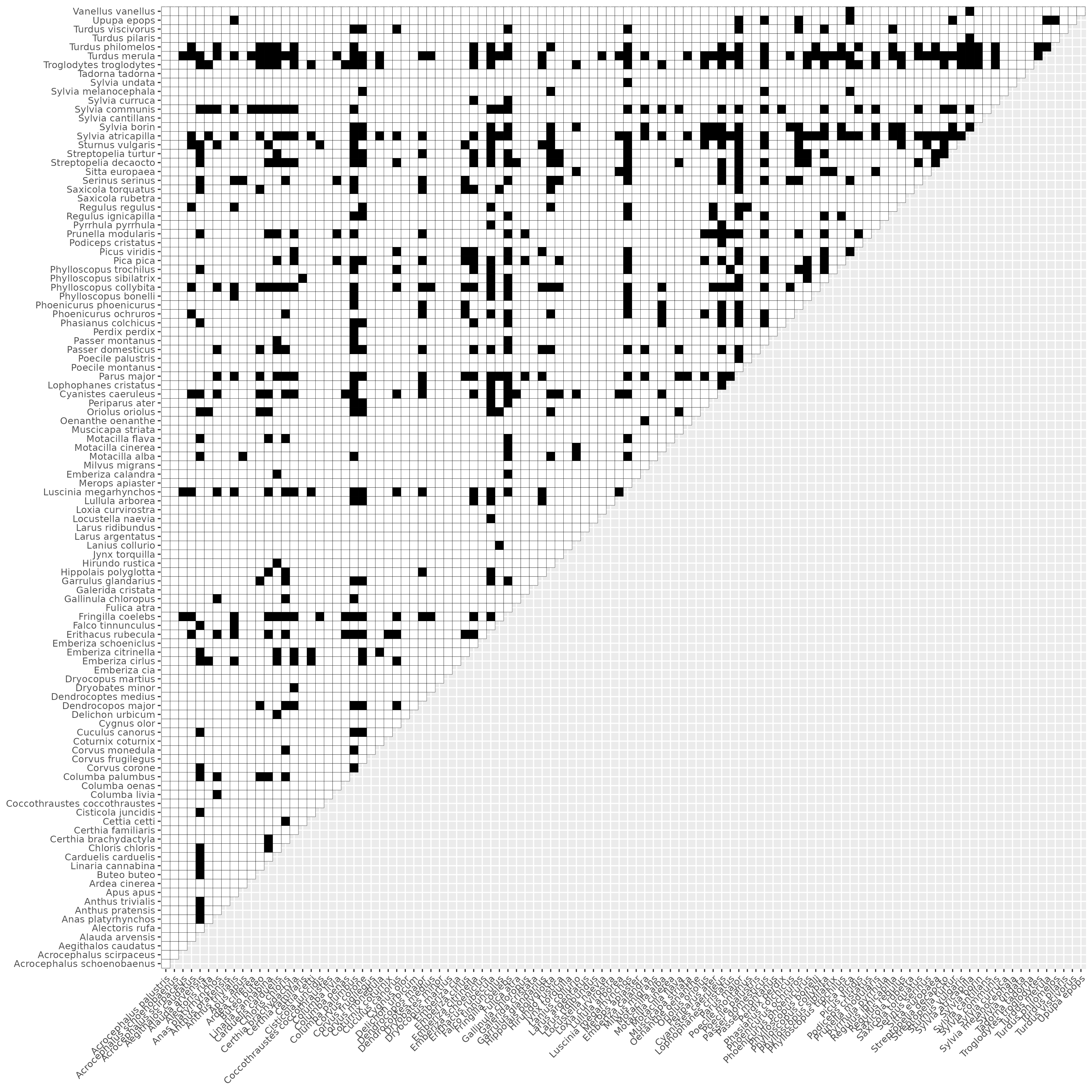
Figure S5A: Matrix of associated species pairs (black cells) in French bird communities.

In addition to validation of theoretical expectations (see main text), several species associations found in our study were in line with current knowledge on species behaviour and interspecific relationships. For instance (Fig. S5A), the Common cuckoo (*Cuculus canorus*) was associated to the European robin (*Erithacus rubecula*), the Cirl bunting (*Emberiza cirlus*), the Common nightingale (*Luscinia megarhynchos*), and the Willow warbler (*Phylloscopus trochilus*), known to be frequent hosts of cuckoo's eggs (Davies and Brooke, 1989; Moksnes and ØSkaft, 1995). The Great spotted woodpecker (*Dendrocopos major*) is associated with the secondary cavity nesters, such as the Common redstart (*Phoenicurus phoenicurus*), the Great tit (*Parus major*), the Eurasian blue tit (*Cyanistes caeruleus*) and the Crested tit (*Lophophanes cristatus*) (van der Hoek et al., 2017).
